# A Generalized Supervised Contrastive Learning Framework for Integrative Multi-omics Prediction Models

**DOI:** 10.1101/2023.11.01.565241

**Authors:** Sen Yang, Shidan Wang, Yiqing Wang, Ruichen Rong, Bo Li, Andrew Y. Koh, Guanghua Xiao, Dajiang Liu, Xiaowei Zhan

## Abstract

Recent technological advances have highlighted the significant impact of the human microbiome and metabolites on physiological conditions. Integrating microbiome and metabolite data has shown promise in predictive capabilities. We developed a new supervised contrastive learning framework, MB-SupCon-cont, that (1) proposes a general contrastive learning framework for continuous outcomes and (2) improves prediction accuracy over models using single omics data. Simulation studies confirmed the improved performance of MB-SupCon-cont, and applied scenarios in type 2 diabetes and high-fat diet studies also showed improved prediction performance. Overall, MB-SupCon-cont is a versatile research tool for multi-omics prediction models.

## 1 Introduction

Integrated modeling approaches that synergistically combine multi-omics data have revolutionized biomedical research, enabling comprehensive interrogation of the complex biological relationships between multiple aspects (e.g., microbial communities and metabolic activities) of human physiology. The human microbiome, a diverse array of living microorganisms residing in a variety of body niches, plays a pivotal role in diseases diagnosis, prognosis, and treatments. [1-3] The metabolome, a complete set of small molecules involved in metabolic processes, impacts the host physiology conditions. [4-6] By utilizing multi omics data, researchers have developed accurate prediction models for host physiological conditions, with translational potential for improved healthcare. [7, 8]

Early work on prediction modeling relied on single-omics data and methods such as elastic net, random forests, support vector machines, and gradient boosting trees. [9-11] However, integrating multi-omics data offers mechanistic insights and can potentially enhance the accuracy of prediction models. For instance, in colorectal cancer, researchers have identified specific bacterial species that produce metabolites linked to an increased risk of the disease. [12] Subsequent mechanistic studies have further elucidated the functions of these pathogenic species through multi-omics analysis. [13] Similar multi-omics approaches, particularly in microbiome and metabolomics, have been applied to other diseases as well. [14-16]

Existing integration models based on multivariate statistics exhibit potential, but recent advances in deep learning, particularly contrastive learning, have shown greater promise. Contrastive learning has improved image classification in computer vision and pretraining language models in natural language processing. [17, 18] For example, self-supervised contrastive learning [17] and supervised contrastive learning [19] have been successful in image recognition. Recently, contrastive learning has also been introduced to multi-omics data analysis. Using supervised contrastive learning, MB-SupCon achieves superior performance on predicting various covariates by integrating microbiome and metabolome data. [20] However, existing contrastive learning frameworks have limitations. Their supervised mechanisms only support the prediction of categorical covariates, while continuous covariates are quite common in biological research.

To address these challenges, we propose MB-SupCon-cont, a generalized contrastive learning framework for both categorical and continuous covariates on multi-omics data. Specifically, we generalize the concept of “similar data pairs” based on the distance of responses between two data points and use it in a generalized contrastive loss. We have demonstrated the superiority of MB-SupCon-cont by predicting multiple covariates in both simulation and two real studies. Additionally, the learned embedding on the representation domain can be used to generate gradually changing clusters according to the response variable, which can be utilized for better visualization purposes. Due to these advantages, MB-SupCon-cont can be used as a general integrative prediction modeling framework in multi-omics studies and can be versatilely helpful for the biomedical research community.

Code availability: https://github.com/ya61sen/MB-SupCon-cont

## 2 Methods

### 2.1 Similar and Dissimilar Data Pairs

Contrastive learning leverages the idea of learning embeddings such that similar data points are brought closer in the embedding space, while dissimilar points are pushed apart. Therefore, how to find these similar and dissimilar data pairs are critical in all types of contractive learning methods.

In self-supervised contrastive learning, we define positive (similar) and negative (dissimilar) data pairs based on whether the data is from a paired sample collected simultaneously from the same subject. For a data pair {***x***_*i*_, ***y***_*j*_}; *i, j ∈* {1,2, …, *n*} where *n* is the batch size.

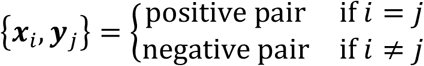

In supervise contrastive learning for categorical response variable, positive and negative data pairs are determined based on whether the data has the same response value. Given a data pair {***x***_*i*_, ***y***_*j*_} and their responses {*r*_*i*_, *r*_*j*_} where *i, j ∈* {1,2, …, *n*}, the positive (or negative) pair is defined as

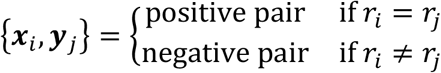

However, when the response variable is continuous for a supervised contrastive learning model, the definition of positive versus negative data pairs defined above is not applicable, as continuous response values are very unlikely to be the same. Instead, similar and dissimilar data pairs are defined based on the distance of response values. For data {***x***_*i*_, ***y***_*j*_} and their continuous responses {*r*_*i*_, *r*_*j*_}, we can define a distance *d*_*ij*_ = |*r*_*i*_ − *r*_*j*_| and use it to define the similarity. A distance nearing 0 indicates a highly similar data pair, while a value approaching its maximum signifies dissimilarity. We will use *d*_*ij*_ in the derived generalized contrastive loss.

### 2.2 Contrastive learning and its loss function

The general idea of contrastive learning is to maximize the similarity of encoded embeddings in the representation domain. To measure the similarity of two vectors, the cosine similarity is the most frequent choice. From a mini-batch point of view, suppose ***x***_*i*_ and ***y***_*j*_ are different omics data vectors obtained from sample *i* and sample *j* (*i, j ∈* {1,2, …, *n*}; *n* is the batch size), respectively,

*f*_*x*_ and *f*_*y*_ are the encoders for each omics data, then the embeddings for ***x***_*i*_ and ***y***_*j*_ are 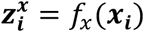 and 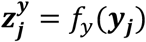. The similarity of embeddings is defined as

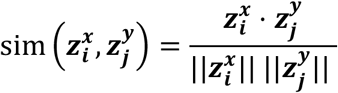

where · denotes the dot product and ‖ · ‖ is the Euclidean norm of a vector.

By anchoring the embedding of one omics data ***z***^***x***^, we define the generalized contrastive loss to be

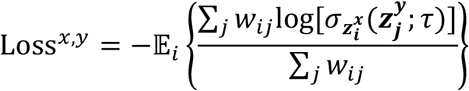

where ***w***_*ij*_ *∈* [0,1] is the weight for sample pair {***x***_*i*_, ***y***_*j*_}, 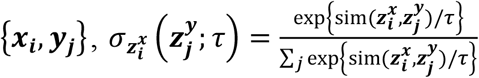 is the SoftMax probability of cosine similarity of scaled embeddings and scaler *τ ∈* ℝ_+_ is the temperature parameter.

By fixing the other embedding ***z***^***y***^ as the anchor, we will get the other piece of loss Loss^*y,x*^. For two omics data, the total contrastive loss is Loss = Loss ^*x, y*^ + Loss^*y, x*^

Rewrite this formula in matrix form:

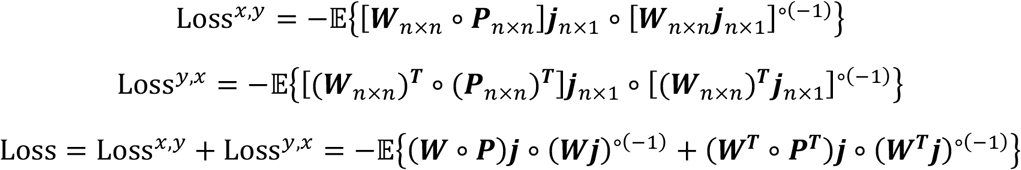

where ○ denotes the Hadamard product (element-wise product) and (·)^°(−1)^ denotes the Hadamard inverse. The Hadamard inverse of matrix ***X*** is defined as 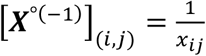. Also, vector *j*_*n*×1_ = [1, 1, …, 1]^T^, matrix ***W***_*n×n*_ is the weight matrix and matrix ***P***_*n×n*_ is the SoftMax probability matrix, with 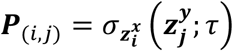.

By this definition, we can use one generalized formula to incorporate self-supervised contrastive loss, supervised contrastive loss on categorical covariates and supervised contrastive loss on continuous covariates. The key lies in the definition of the weight matrix ***W***, which encapsulates the information of both similar and dissimilar pairs.

First, for self-supervised contrastive loss, we have ***W***_n×n_ = ***I***_n×n_ where ***I*** is an identity matrix.

Second, for supervised contrastive loss on categorical covariates, ***W*** is defined as

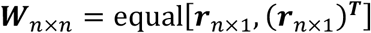

where ***r*** is the response vector and equality function equal(·) is a true/false function to check the element-wise equalities. Equivalently,

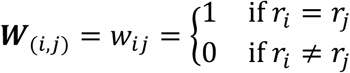

Third, for supervised contrastive loss on continuous covariates, we propose three methods to define the weight matrix ***W***. The underlying principle is to assign a weight to a data pair based on the distance of their responses, with the weight being negatively correlated to the response distance. Namely, the larger the distance is, the smaller the weight is and vice versa.

Suppose *r* _*i*_, *r*_*j*_ are the responses for a data pair {***x***_*i*_, ***y***_*j*_}, the norm-1 distance is *d* _*i j*_ = | *r*_*i*_ − *r* _*j*_ |. Accordingly, the three weights are defined as follows.

(1)Linear weights

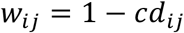

where the scale 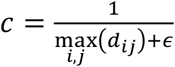 and a small positive constant ϵ is to avoid zero denominator.

(2)Exponential weights

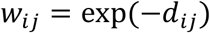

(3)Negative-log weights

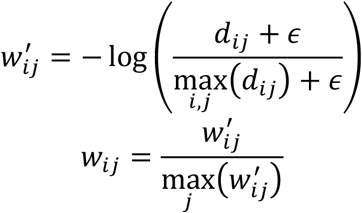

where ϵ is a small positive constant to avoid zero values. Here, we further scale 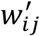 to contain the weights within [0,1].

In the rest of the article, we will use all three methods to train the model and compare the performance between each other. In general, we believe there is hardly one universally ultimate method for any data. The best method depends on the distribution of the responses and the data. We should consider the choice of the weighting method case by case.

### 2.3 MB-SupCon-cont Framework

The initial stage of the MB-SupCon-cont framework is data collection (**Figure 1**). For example, we can collect paired omics data along with patients’ phenotype information. The next step is contrastive learning. Here we train a supervised contrastive learning model and obtain encoders *f*_*x*_ and *f* _*y*_. Note that the generalized contrastive loss should be employed in this context to accommodate various types of covariate data. In the last step, prediction heads (either classifiers or regressors) are utilized on the embeddings. When making new predictions, embeddings from either microbiome data or other omics data should be chosen based on their performance during the training phase. Additionally, a unique trend related to the covariates can be visualized in the lower-dimensional space.

**Figure 1.**
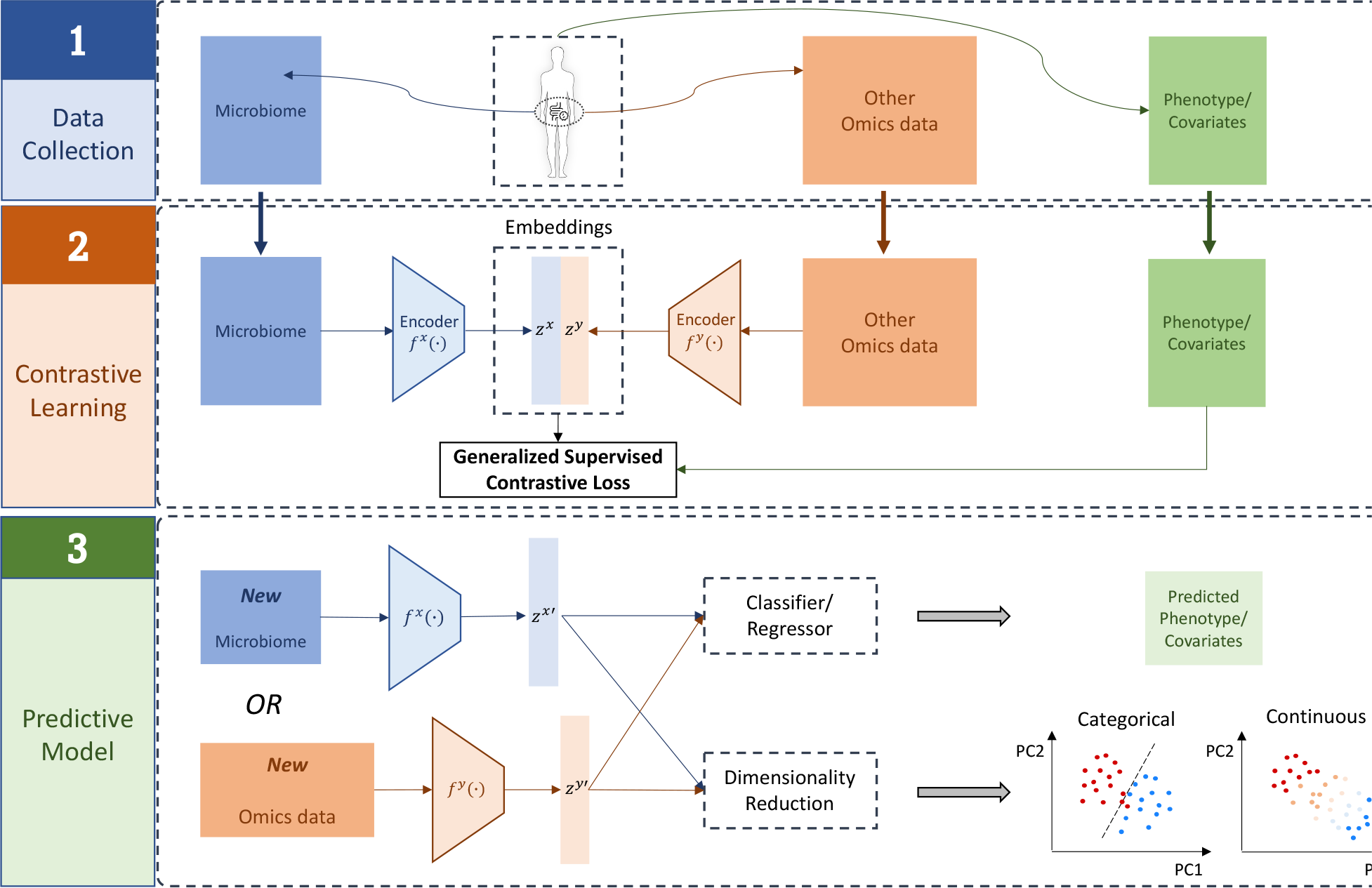
Framework of MB-SupCon-cont. Step 1: collect paired microbiome and other related omics data, as well as phenotype information from the patients. Step 2: train a supervised contrastive learning model using generalized contrastive loss. Step 3: after the model is trained, given either new microbiome data or other omics data, we can improve the prediction of phenotypes and achieve better visualization.

## 3 Results

### 3.1 Simulation study

To benchmark the efficacy of MB-SupCon-cont, we designed a simulation study. The simulation data was generated by Reverse PCA, a technique we developed and detailed in **Supplementary Appendix A**. It reconstructed the original omics data from its lower-dimensional representation. The choice of Reverse PCA was grounded in its simplicity and ease of implementation, making it an accessible yet robust tool for our analysis. It is assumed that for two sets of omics data, a representation space exists wherein the corresponding features from both datasets are correlated. Reverse PCA is used to reconstruct a reduced-rank approximation of the original data based on their principal components. This approach guaranteed that the vital information contained within the embeddings is preserved through the reconstruction process. In this study, we assume the generated embeddings in the representation space are corresponding to the highest-variance principal components. Furthermore, the response variable is linked to these embeddings.

To showcase the efficacy and benefits of MB-SupCon-cont, the prediction on simulated original data is compared to the prediction on learned embeddings. The prediction heads under consideration are: (1) Elastic net regression; (2) Support vector regression (SVR); (3) Random forest regression (RF); (4) Extreme boosting regression (XGBoost).

We generate 1000 paired samples and set the dimension of original data to be *D*_*x*_ = 100, *D*_*y*_ = 500. The response variable is generated using reverse PCA. Specifically, the coefficients were generated from a normal distribution *N* (0, 25), and the generated response is scaled to 0 − 100 assuming it contains Age information. A set of *μ*_*ρ*_ *∈ {*0.4, 0.6, 0.8*}* is considered for the average correlation of features, and a set of *d*_*e*_ *∈ {*10, 20, 40*}* is tested for the dimension of embeddings (**Figure 2 and Supplementary Figure 1 and 2**).

**Figure 2.**
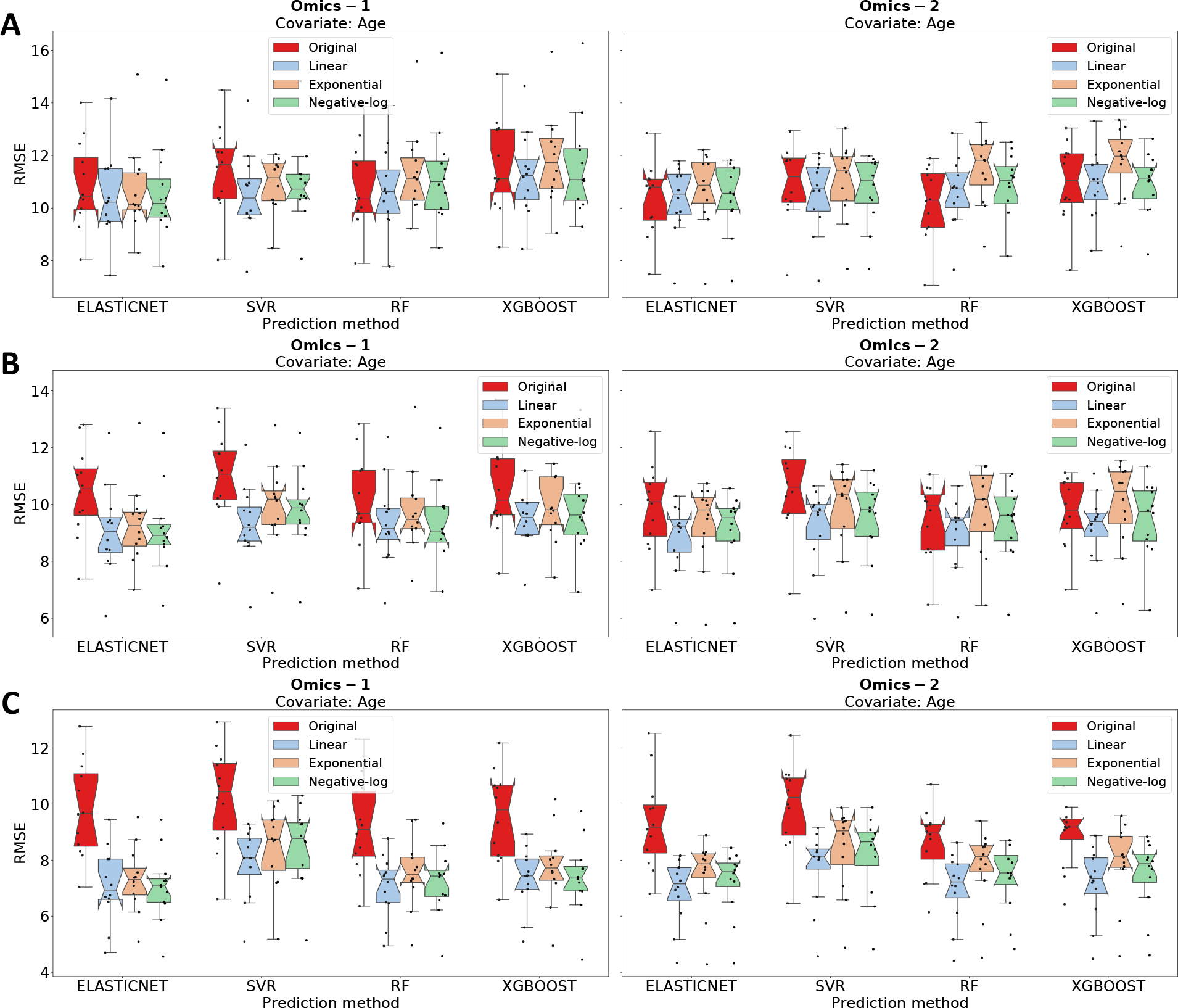
Boxplots of prediction RMSEs on testing data (15%) of simulation study from 12 random replicates when the dimension of embeddings is **20**. Panel A: *μ*_*ρ*_ = 0.4; Panel B: *μ*_*ρ*_ = 0.6; Panel C: *μ*_*ρ*_ = 0.8. Red boxes represent the prediction on original data. The light blue, orange, and green boxes represent the prediction on embedding learning by using linear, exponential, and negative-log weighting methods.

Our simulation demonstrated the superiority of MB-SupCon-cont in predicting continuous covariates. Specifically, as the average correlation of embedding features rises, the advantage of MB-SupCon-cont grows correspondingly. At *μ*_*ρ*_ = 0.8, the benefit of using MB-SupCon-cont is most pronounced. In most scenarios, the light-colored bars (prediction on embeddings) are significantly lower than the red bar (prediction on original data). Additionally, the use of linear weights results in marginally better performance than the other two weighting methods when predicting the simulated “Age” variable. This illustrates that MB-SupCon-cont can enhance prediction performance based on embeddings, and this enhancement intensifies with the average correlation coefficient of embeddings.

### 3.2 Application – Type 2 Diabetes (T2D) Study

We benchmarked MB-SupCon-cont to paired microbiome and metabolome data [21]. This study enrolled 720 samples of prediabetes patients. Three continuous covariates that are all associated with the disease status of diabetes patients were considered for prediction, which are Age, Body Mass Index (BMI) and Steady State Plasma Glucose (SSPG) level.

To assess the prediction performance, we employed the Root-Mean-Square Error (RMSE) as the evaluation metric. For a comparative perspective, predictions were also made using the original microbiome and metabolome data. The dataset was partitioned into a training set (70%), a validation set (15%), and a testing set (15%). The dimensions of the microbiome data are 720 *×* 96, for metabolome data, they are 720 *×* 724. We executed the analysis 12 times, each with unique random seeds for the training-validation-testing splits, and recorded the RMSE for all prediction methodologies during each run.

In **Table 1**, the mean RMSE values averaged over the 12 iterations, are contrasted across the various prediction methods. For predictions based on MB-SupCon-cont embed-dings, only results by using random forest (MB-SupCon-cont + RF) are presented as it has the best overall performance. Prediction results of other predictors can be found in **Supplementary Appendix B.1**. Among the three weighting methods, MB-SupCon-cont using linear weights has the best performance on all three covariates. Furthermore, predictions based on metabolome embeddings using the linear weighting method outperform all the other prediction methods as it has the lowest RMSE. By employing MB-SupCon-cont, the predictions of all three continuous covariates are improved by a large margin. The best average RMSE for predicting Age is 4.0593, for predicting BMI is 1.4826, and for predicting SSPG is 18.4474. A boxplot comparison of all prediction methods for predicting three covariates is also presented in **Figure 3**. The light blue box (MB-SupCon using linear weighting method) in the metabolome side always reaches the lowest RMSE, holding a consistent result with **Table 1**.

**Table 1.** Average RMSEs on testing data (T2D study) for predicting Age, BMI and SSPG by different predictors based on original omics data and MB-SupCon-cont embeddings using three weighting methods (italic in the table). For predictions based on embeddings, only results by using random forest (MB-SupCon-cont + RF) is presented.

| **Microbiome** | **Original** |  |  |  | **MB-SupCon-cont + RF** |  |  |
| --- | --- | --- | --- | --- | --- | --- | --- |
|  | ElasticNet | SVR | RF | XGBoost | *Linear* | *Exponential* | *Negative-log* |
| **Age** | 9.4462 | 9.6207 | 7.8069 | 8.1214 | 7.6823 | 8.3237 | 7.8280 |
| **BMI** | 3.8306 | 3.5244 | 2.9652 | 3.1530 | 3.1830 | 3.5324 | 3.4105 |
| **SSPG** | 48.1111 | 59.0257 | 41.9548 | 44.1595 | 45.0804 | 54.2055 | 45.4351 |

| **Metabolome** | **Original** |  |  |  | **MB-SupCon-cont + RF** |  |  |
| --- | --- | --- | --- | --- | --- | --- | --- |
|  | ElasticNet | SVR | RF | XGBoost | *Linear* | *Exponential* | *Negative-log* |
| **Age** | 6.3917 | 8.8224 | 5.6916 | 5.7289 | 4.0593 | 5.0115 | 5.3216 |
| **BMI** | 2.8708 | 2.8792 | 2.1848 | 2.1154 | 1.4826 | 1.7599 | 1.6190 |
| **SSPG** | 41.7313 | 59.3345 | 34.4950 | 33.5242 | 18.4474 | 46.9629 | 23.4287 |

**Figure 3.**
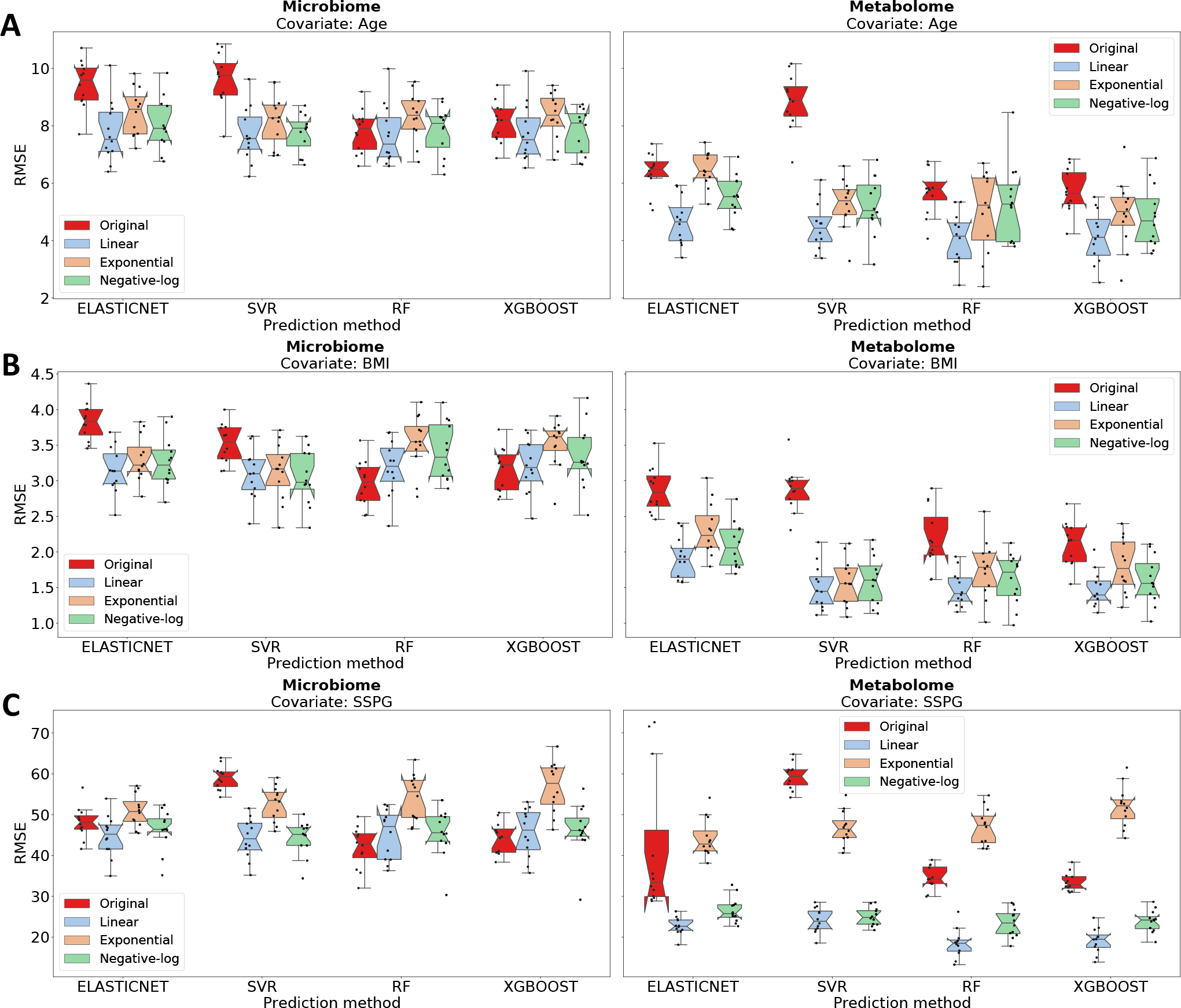
Boxplots of prediction RMSEs on testing data from 12 random training-validation-testing splits, by using different methods for three continuous covariates (T2D study). Panel A: Age; Panel B: BMI; Panel C: SSPG. Red boxes represent the prediction on original data. The light blue, orange and green boxes represent the prediction on embeddings learning by using linear, exponential, and negative-log weighting methods.

For comparison, principal component analysis (PCA) was performed on both (1) the original microbiome and metabolome data, and (2) microbiome and metabolome embeddings of MB-SupCon-cont using linear weighting method. From **Supplementary Figure 3** (metabolome) and **Supplementary Figure 4** (microbiome), both metabolome and microbiome embeddings (second rows in each figure) have shown a much more distinctive varying trend on the lower-dimensional space than the original metabolome and microbiome data (first rows in each figure). This shows that MB-SupCon-cont embeddings provide accurate covariate-related representations of the original data and are beneficial for visualization.

### 3.3 Application – High-Fat Diet (HFD) Study

To further validate the superiority of MB-SupCon-cont in predicting continuous covariates, we applied it to another independent multi-omics murine model studying the effect of a high-fat diet on the microbial and metabolic changes in mice [22]. This model utilized 434 paired microbiome and metabolome data for integration. In contrast to the previous T2D study, this HFD study presented a considerably larger dimensionality in the paired omics data, with the microbiome data dimension being 434 × 913 and the metabolome data dimension being 434 × 11978. We still split the dataset into training (70%), validation (15%), and testing data (15%). The analysis was conducted 12 times using different random seeds. Covariate Age and Weight are selected as the prediction tasks.

**Supplementary Table 1** and **Supplementary Figure 5** have shown the comparison of prediction RMSEs on testing data (15% of all data) by different methods. Similarly as T2D study, prediction based on metabolome embeddings from MB-SupCon-cont using linear weighting method reached the best performance (lowest average RMSE) for both Age and Weight predictions. The most accurate predictions were obtained using metabolome embeddings, resulting in a RMSE of 7.1592 for Age and 6.4430 for Weight. Meanwhile, the color variations corresponding to different values of the variable are clearly distinguishable in the lower-dimensional PCA scatter plots (PC2 vs. PC1) of metabolome and microbiome embeddings (**Supplementary Figure 6** and **Supplementary Figure 7**). This study confirms the stable and consistent performance of MB-SupCon-cont.

## 4 Discussion

We develop a novel supervised contrastive learning framework. We design a new weighting scheme that extends the general supervised contrastive learning framework. It can integrate multi-omics datasets by deriving omic-specific embeddings and achieve three advantages: (1) enhanced predictive performance (lower RMSE): The model demonstrates superior prediction performance, enabling more accurate and reliable analysis of multi-omics data. (2) Advanced representation learning: Employing embeddings as a means of representation learning facilitates the extraction of meaningful and comprehensive information from the data. (3) Enhanced visualization on lower-dimensional space: The model yields enhanced visualization, unveiling a smooth yet distinguishable color trend in PCA scatter plots for continuous variables. We have demonstrated its performance through extensive simulation and real data analysis.

Given the outstanding performance of MB-SupCon and MB-SupCon-cont using supervised contrastive learning methods, an essential question arises: What criteria should we consider when selecting datasets for integration? It’s clear that not every pair of data can be effectively integrated. The rationale for choosing datasets for MB-SupCon and MB-SupCon-cont is related to the concept of “informativeness” [23]. An informative pair of datasets should neither be “easy positives” (too similar to offer more information than a single dataset) nor “easy negatives” (so unrelated that they are entirely different from each other). Based on this idea, in biological studies, the chosen datasets should be biologically connected yet offer diverse perspectives on a system.

Another aspect to consider with MB-SupCon-cont is the selection of weighting methods. While we introduce three weighting methods, linear weights appear to outperform in most studies. However, we posit that the choice of weighting methods largely depends on the distribution of the original data and the responses. A potential avenue for future research is to determine the optimal weighting methods for supervised contrastive loss. Through simulation and real data analysis, a more universally applicable guideline is sought.

In addition, while we demonstrated the utilities of MB-SupCon-cont in integrative microbiome and metabolomics datasets, this work can be extended to other omics types. For example, integrating host genetics, genomics, epigenetics, and proteomics, along with microbiome and metabolomics is also feasible. For example, we discovered that proteomics measurements can improve microbiome-based prediction models in the iHMP IBD datasets. Thus, our proposed framework has the potential to improve integrative data analysis in broader settings.

In conclusion, MB-SupCon-cont, coupled with encoder-based neural networks, possesses a distinct advantage in the approximation of non-linear functions and the modeling of high-dimensional data. The versatility of the MB-SupCon-cont framework makes it applicable in a wide range of multi-omics scenarios, ultimately enhancing the efficacy of microbiome-based prediction models.

