## Supplementary Materials for "A Generalized Supervised Contrastive Learning Framework for Integrative Multi-omics Prediction Models"

**Supplementary Figure 1.** Boxplots of prediction root-mean-square errors (RMSEs) on testing data (15%) of simulation study from 12 random replicates when the dimension of embeddings is **10**. Panel A: $\mu_\rho = 0.4$; Panel B: $\mu_\rho = 0.6$; Panel C: $\mu_\rho = 0.8$. Red boxes represent the prediction on original data. The light blue, orange and green boxes represent the prediction on embeddings learning by using linear, exponential, and negative-log weighting methods.

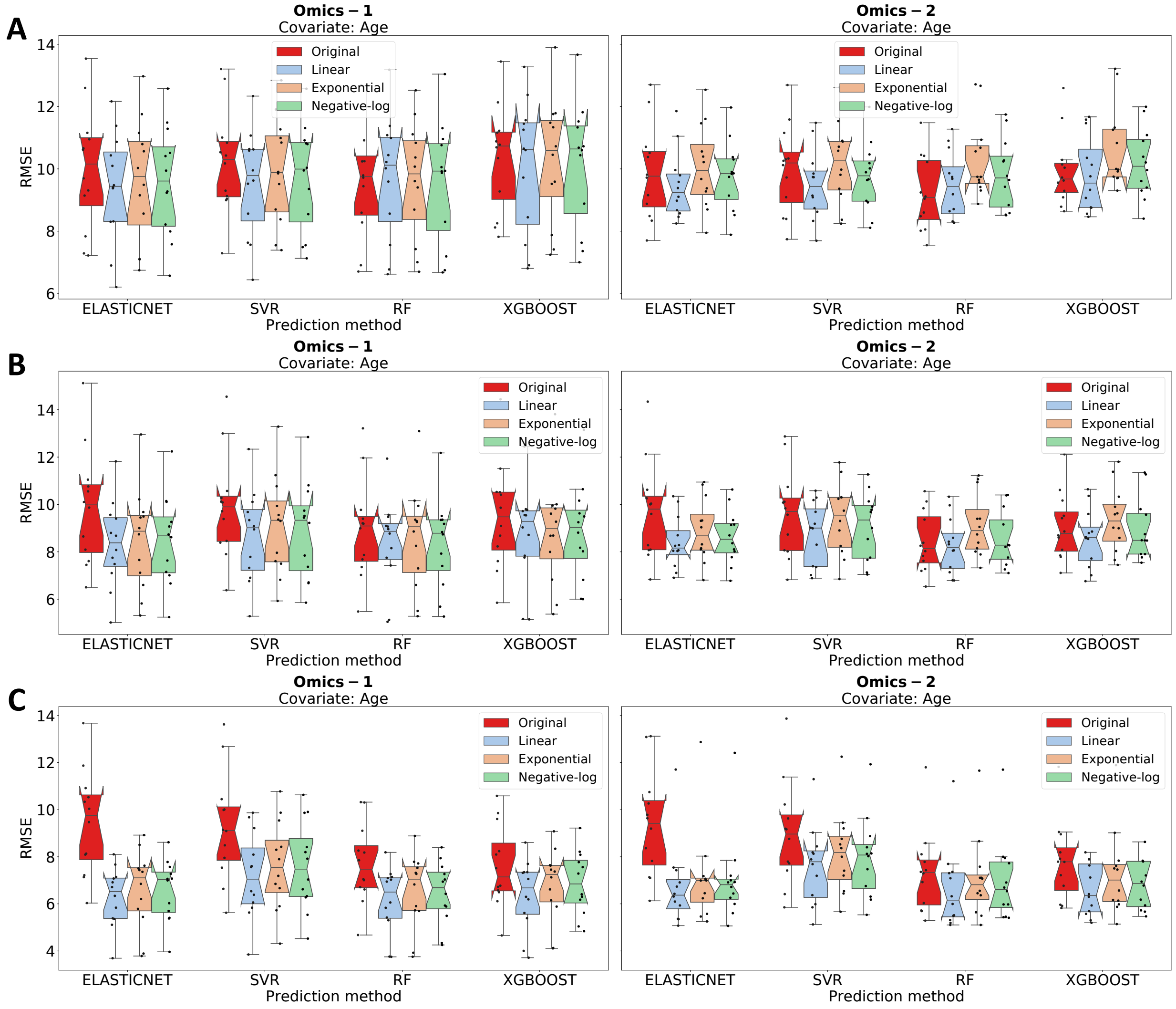

**Supplementary Figure 2.** Boxplots of prediction RMSEs on testing data (15%) of simulation study from 12 random replicates when the dimension of embeddings is **40**. Panel A: μρ = 0.4; Panel B: μρ = 0.6; Panel C: μρ = 0.8. Red boxes represent the prediction on original data. The light blue, orange and green boxes represent the prediction on embeddings learning by using linear, exponential, and negative-log weighting methods.

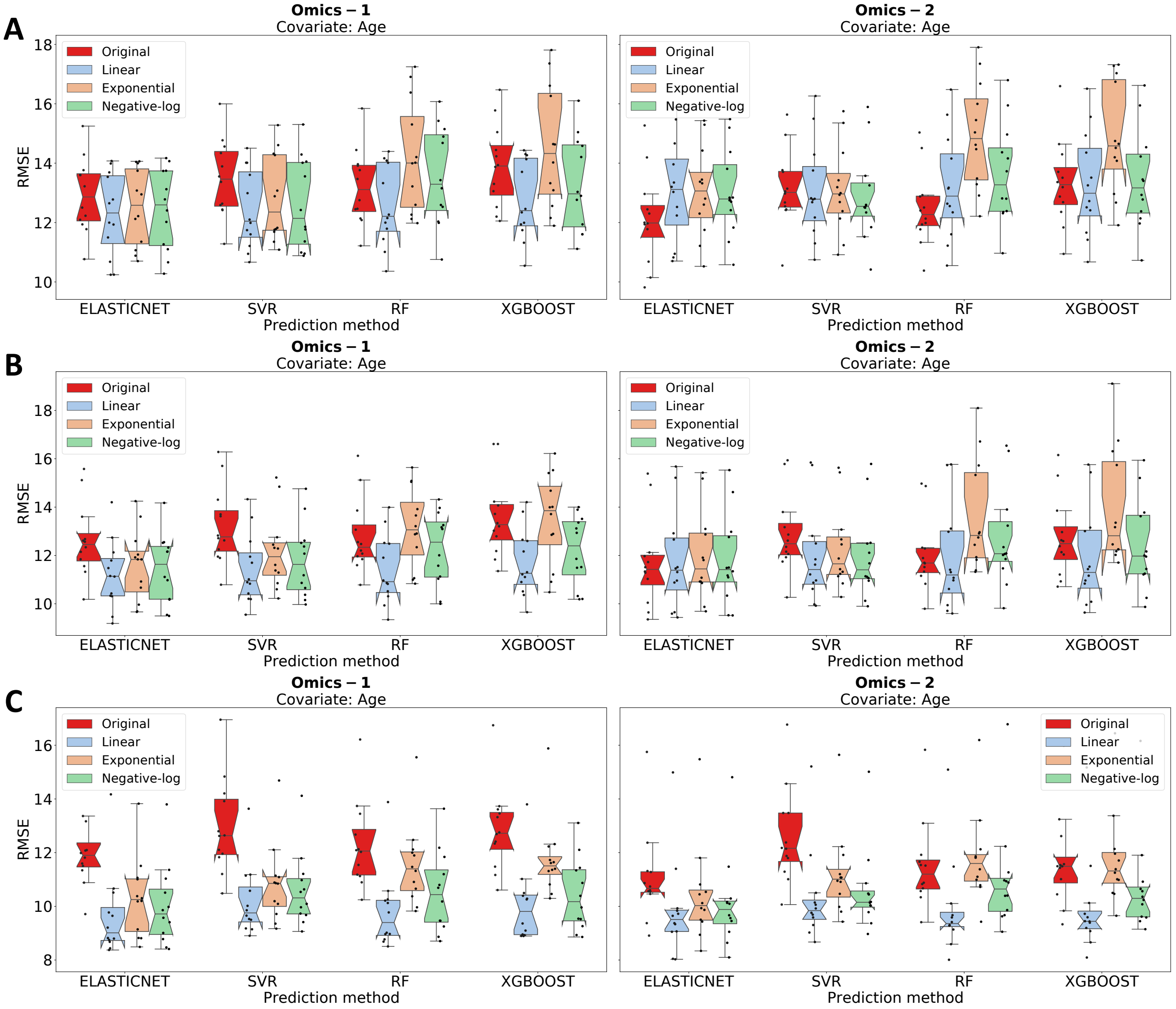

**Supplementary Figure 3.** PCA scatter plots (PC2 vs. PC1) on original metabolome data vs. MB-SupCon-cont metabolome embeddings for covariates Age, BMI, and SSPG (T2D study). First row: original data; Second row: embeddings.

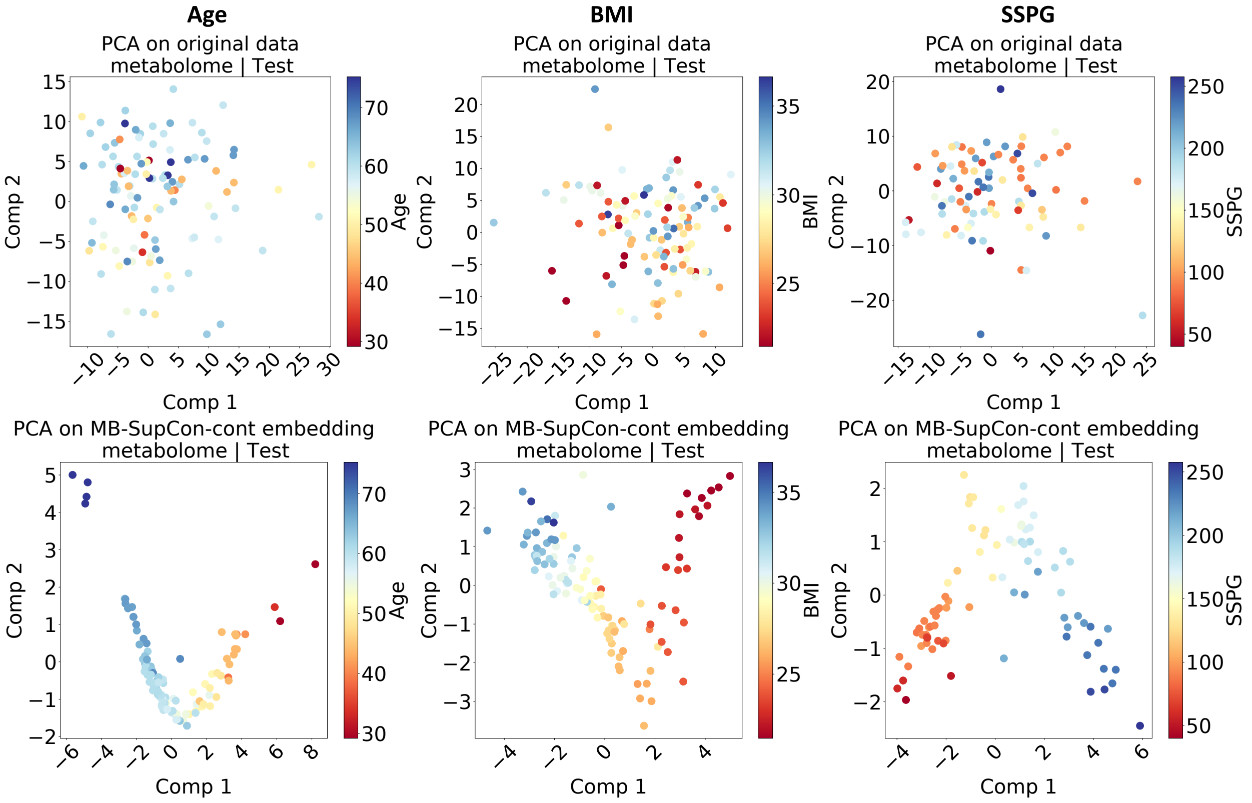

**Supplementary Figure 4.** PCA scatter plots (PC2 vs. PC1) on original microbiome data vs. MB-SupCon-cont microbiome embeddings for covariates Age, BMI and SSPG (T2D study). First row: original data; Second row: embeddings.

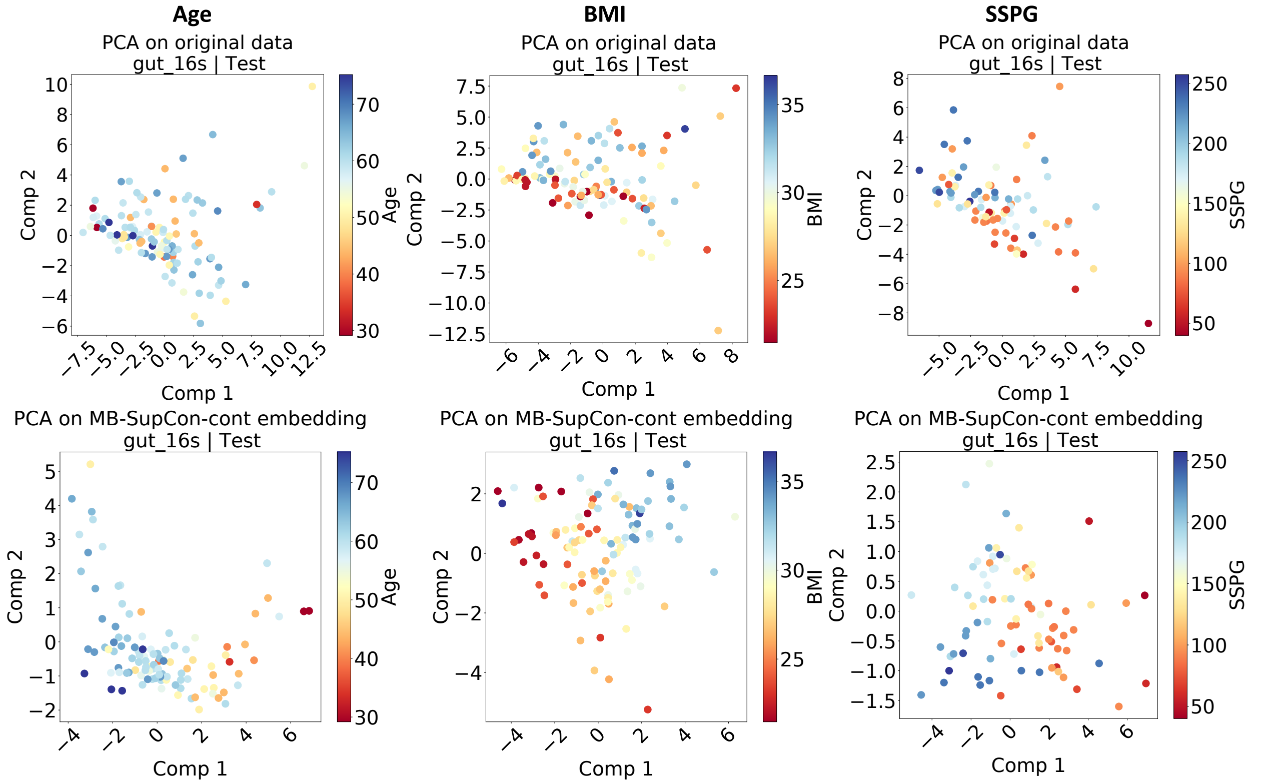

**Supplementary Figure 5.** Boxplots of prediction RMSEs on testing data from 12 random training-validation-testing splits, by using different methods for two continuous covariates (HFD study). Panel A: Age; Panel B: Weight. Red boxes represent the prediction on original data. The light blue, orange and green boxes represent the prediction on embeddings learning by using linear, exponential, and negative-log weighting methods.

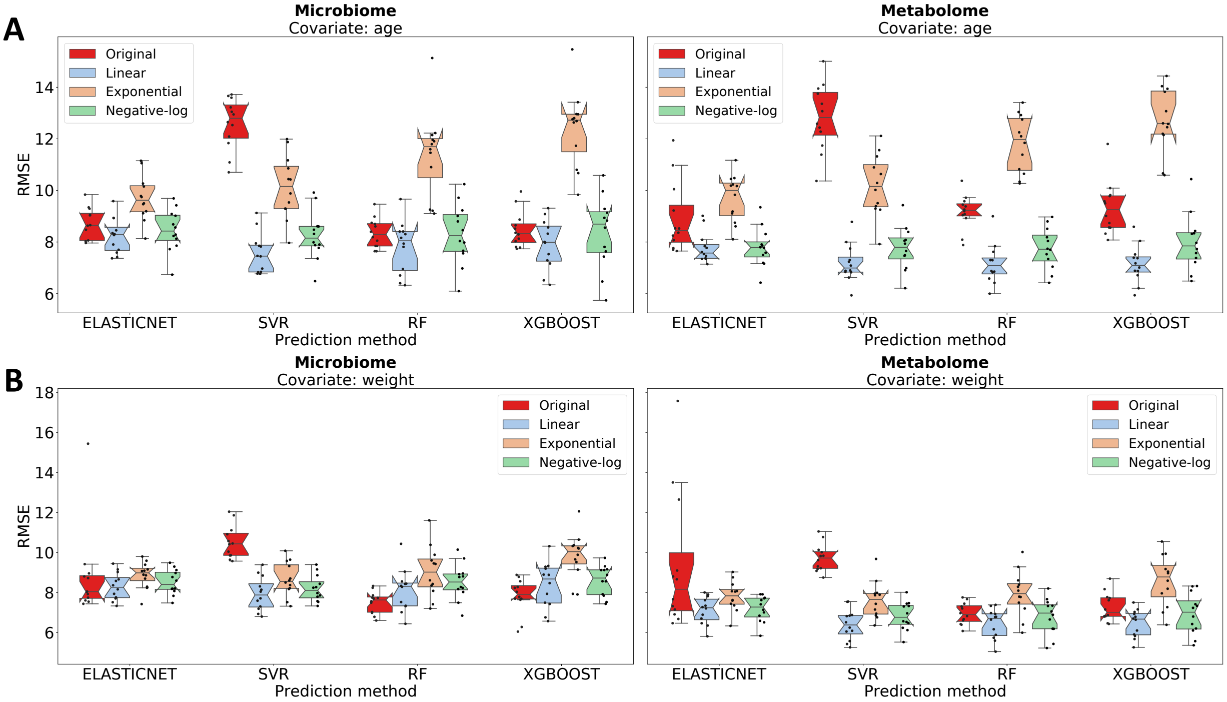

**Supplementary Figure 6.** PCA scatter plots (PC2 vs. PC1) on original metabolome data vs. MB-SupCon-cont metabolome embeddings (HFD study) for covariates Age and Weight. First row: original data; Second row: embeddings.

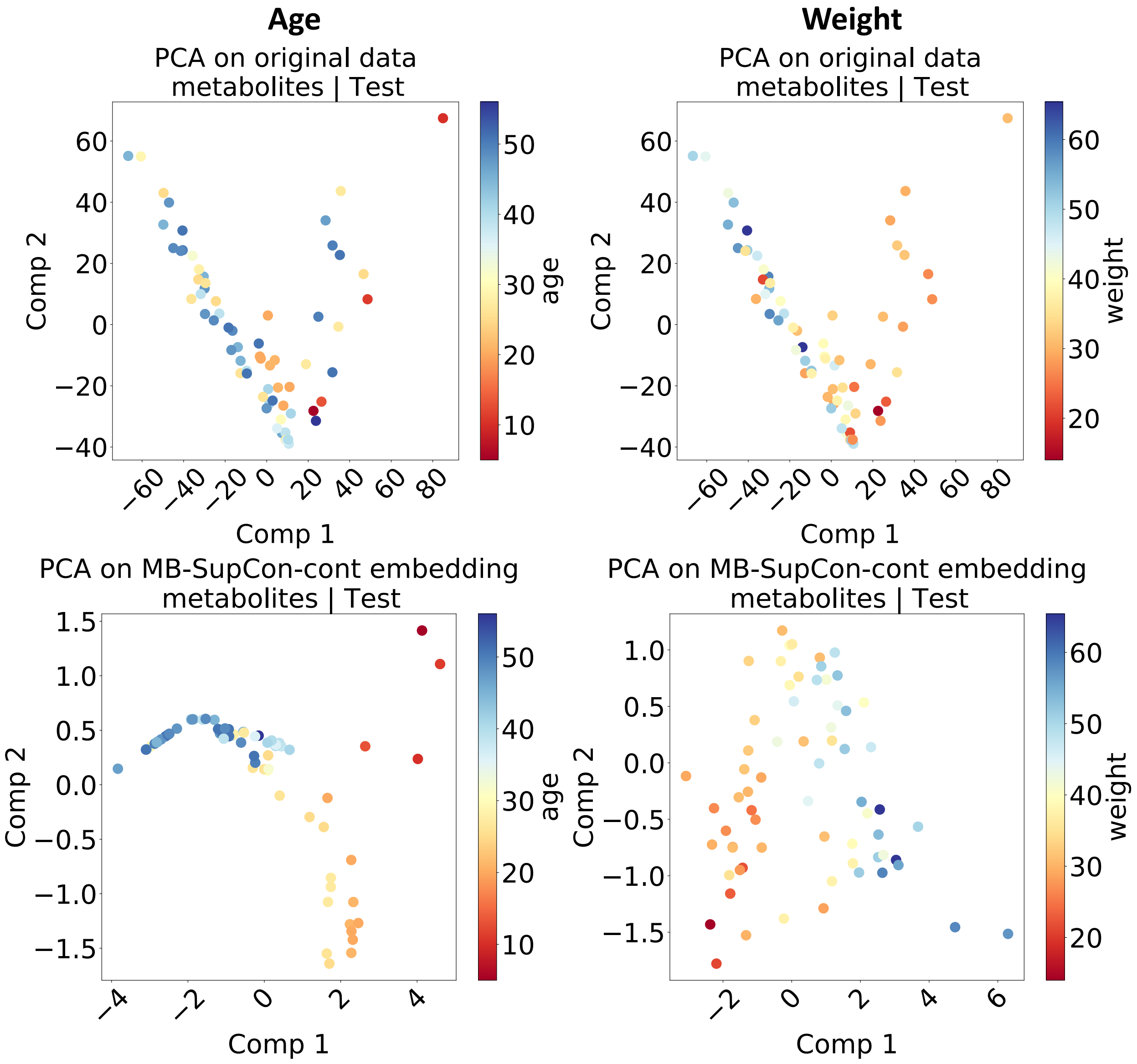

**Supplementary Figure 7.** PCA scatter plots (PC2 vs. PC1) on original microbiome data vs. MB-SupCon-cont microbiome embeddings (HFD study) for covariates Age and Weight. First row: original data; Second row: embeddings.

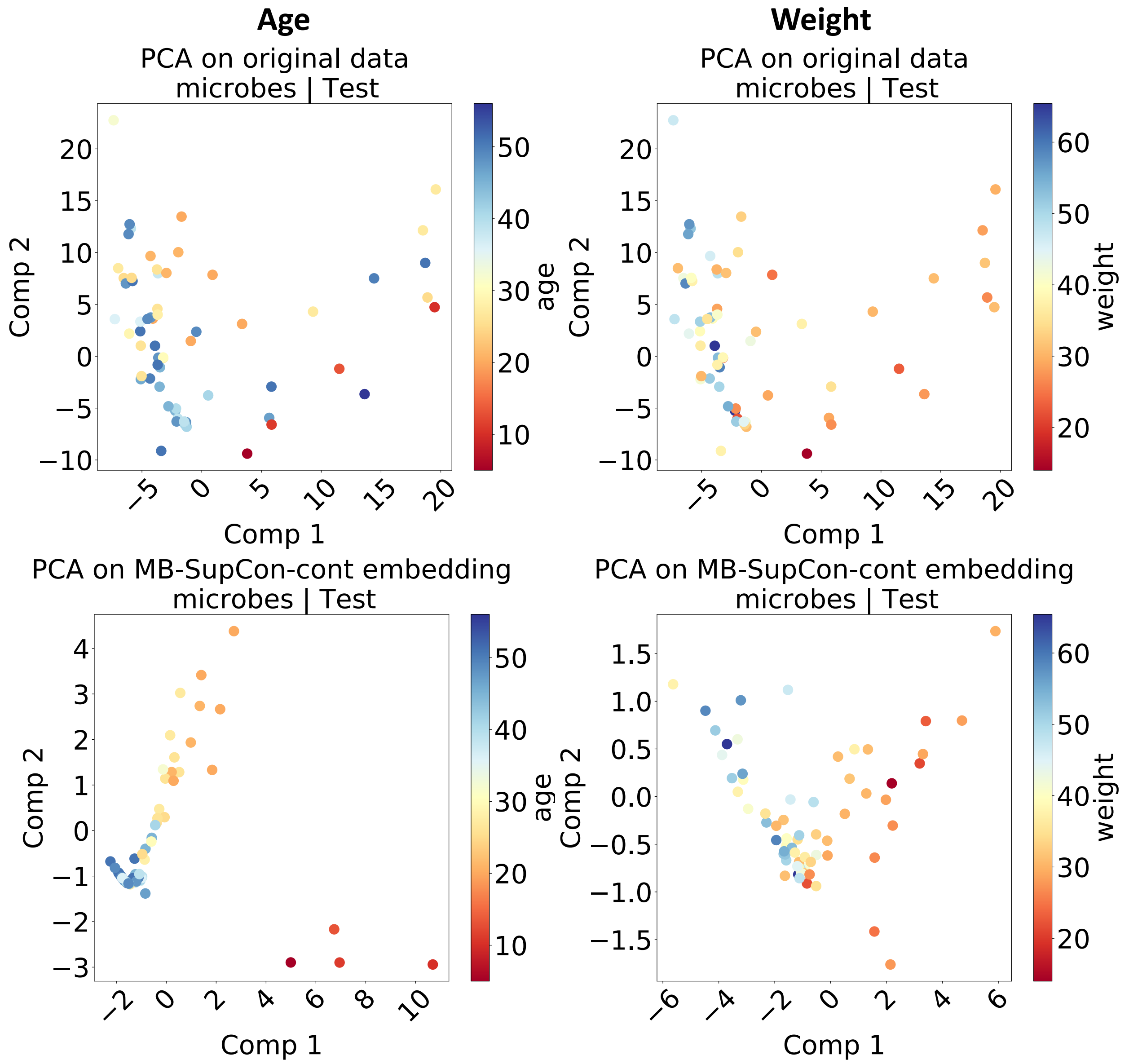

**Supplementary Table 1.** Average RMSEs on testing data (HFD study) for predicting Age and Weight by different prediction heads based on original omics data and MB-SupCon-cont embeddings using three weighting methods (italic in the table). For predictions based on embeddings, only results by using random forest (MB-SupCon-cont + RF) are presented.

| **Microbiome** | **Original** | | | | **MB-SupCon-cont + RF** | | |
|---|---|---|---|---|---|---|---|
| | ElasticNet | SVR | RF | XGBoost | *Linear* | *Exponential* | *Negative-log* |
| **Age** | 8.6952 | 12.5909 | 8.3439 | 8.4767 | 7.8090 | 11.3919 | 8.3139 |
| **Weight** | 8.8105 | 10.5297 | 7.5095 | 7.8293 | 8.1008 | 9.0413 | 8.5323 |
| **Metabolome** | **Original** | | | | **MB-SupCon-cont + RF** | | |
| | ElasticNet | SVR | RF | XGBoost | *Linear* | *Exponential* | *Negative-log* |
| **Age** | 8.9193 | 12.8361 | 9.1652 | 9.3198 | 7.1592 | 11.8144 | 7.7460 |
| **Weight** | 9.3766 | 9.7355 | 6.9497 | 7.2810 | 6.4430 | 7.9505 | 6.8123 |

### Appendix A. Reverse PCA algorithm for simulation study

**Algorithm 1** Simulation via Reverse PCA

Denote the dimension of original data as $D_x, D_y$, the dimension of embedding as $d_e$, the average correlation of features as $\mu_\rho$ and the standard deviation of correlation as $\sigma_\rho^2$.

**Step 1: Generate embeddings.**

1: **for** $k = 1, 2, ..., d_e$ **do**

2: Generate embeddings $\mathbf{z_x}^{(k)}, \mathbf{z_y}^{(k)}$ for omics data $\mathbf{x}$ and $\mathbf{y}$, respectively, s.t.,

$$(\mathbf{z_x}^{(k)}, \mathbf{z_y}^{(k)}) \sim \mathrm{MVN}(\boldsymbol{\mu}^{(k)}, \Sigma^{(k)}) \quad \text{with } \Sigma^{(k)} = \begin{pmatrix} 1 & \rho^{(k)} \\ \rho^{(k)} & 1 \end{pmatrix}$$

where the correlation of corresponding embedding features $\rho^{(k)} \sim N(\mu_\rho, \sigma_\rho^2)$ and $\Sigma^{(k)}$ needs to be positive semi-definite.

3: **end for**

4: **return** Embedding matrices

$$\mathbf{Z_x} = [\mathbf{z_x}^{(1)}, \mathbf{z_x}^{(2)}, ..., \mathbf{z_x}^{(d_e)}],\ \mathbf{Z_y} = [\mathbf{z_y}^{(1)}, \mathbf{z_y}^{(2)}, ..., \mathbf{z_y}^{(d_e)}].$$

**Step 2: Generate original data.**

1: Generate diagonal matrices with ordered explained variances for each principal component

$$\mathbf{V_x} = \mathrm{diag}(v_{(1)}, v_{(2)}, ..., v_{(D_x)}), \quad \mathbf{V_y} = \mathrm{diag}(v'_{(1)}, v'_{(2)}, ..., v'_{(D_y)}).$$

where the unordered explained variances before exaggeration are generated from an exponential distribution: $v_i^{1/c} \sim \mathrm{Exp}(\lambda)$ for $i = 1, 2, ..., D_x$. with scale parameter $\lambda = 1$ and an exaggeration factor $c = 3$, and similarly $v_j'^{1/3} \sim \mathrm{Exp}(1)$ for $j = 1, 2, ..., D_y$.

2: Generate orthogonal matrices $\mathbf{Q}_{\mathbf{x}(D_x \times D_x)}$ and $\mathbf{Q}_{\mathbf{y}(D_y \times D_y)}$ by applying QR decompositions to normally distributed square matrices.

3: Loading matrices are:

$$\mathbf{L_x} = \mathbf{Q_x}\sqrt{\mathbf{V_x}}, \quad \mathbf{L_y} = \mathbf{Q_y}\sqrt{\mathbf{V_y}}.$$

4: Retain only the first $d_e$ principal components and discard the remaining lower-variance principal components:

$$\mathbf{L_x}^{(\text{reduced})} = \mathbf{L_x}[:, 1 : d_m], \mathbf{L_y}^{(\text{reduced})} = \mathbf{L_y}[:, 1 : d_m]$$

5: **return** Reduced-rank approximations of the standardized original omics data:

$$\mathbf{X} = \mathbf{Z_x}[\mathbf{L_x}^{(\text{reduced})}]^T, \quad \mathbf{Y} = \mathbf{Z_y}[\mathbf{L_y}^{(\text{reduced})}]^T.$$

**Step 3: Generate a response variable.**

1: Generate coefficient vectors $\boldsymbol{\beta}_{\boldsymbol{x}(d_e \times 1)}, \boldsymbol{\beta}_{\boldsymbol{y}(d_e \times 1)}$.
2: Suppose the response is quadratically associated with both omics data embeddings.
3: **return** The response variable

$$\mathbf{r} = \mathbf{Z}_{\mathbf{x}}\text{diag}(\boldsymbol{\beta}_{\boldsymbol{x}}) + \mathbf{Z}_{\mathbf{y}}\text{diag}(\boldsymbol{\beta}_{\boldsymbol{y}}) + \mathbf{Z}_{\mathbf{x}} \circ \mathbf{Z}_{\mathbf{x}}\text{diag}(\boldsymbol{\beta}_{\boldsymbol{x}}) + \mathbf{Z}_{\mathbf{y}} \circ \mathbf{Z}_{\mathbf{y}}\text{diag}(\boldsymbol{\beta}_{\boldsymbol{y}}) + \\ \mathbf{Z}_{\mathbf{x}} \circ \mathbf{Z}_{\mathbf{y}}\text{diag}(\boldsymbol{\beta}_{\boldsymbol{x}})\text{diag}(\boldsymbol{\beta}_{\boldsymbol{y}}) + \epsilon.$$

where $\circ$ denotes the Hadamard product and the random noise $\epsilon \sim N(0, 1)$.

**Step 4: MB-SupCon-cont.**

1: Input $(\mathbf{X}, \mathbf{Y})$ to MB-SupCon-cont and make predictions on $\mathbf{r}$.

### Appendix B – Detailed prediction results for real studies

#### B.1. Prediction results for T2D study

In T2D study, the mean RMSEs for predictions based on original omics data and on MB-SupCon-cont embeddings are listed here. The prediction results are summarized for covariates Age and Weight; omics data of microbiome and metabolome; and predictors ElasticNet, SVR, RF and XGBoost. The dimension of embedding is set to be 10.

1. **Covariate**: Age

   **Omics data**: microbiome

Table B.1: Average RMSEs on testing data for predicting Age based on microbiome data and microbiome embeddings from MB-SupCon-cont using three weighting methods (T2D study)

| | **Original** | **MB-SupCon-cont** | | |
|---|---|---|---|---|
| | | Embedding: Linear | Embedding: Exponential | Embedding: Negative-log |
| **ElasticNet** | 9.4462 | 7.7745 | 8.4734 | 8.0504 |
| **SVR** | 9.6207 | 7.6942 | 8.2101 | 7.7500 |
| **RF** | 7.8069 | 7.6823 | 8.3237 | 7.8280 |
| **XGBoost** | 8.1214 | 7.7428 | 8.3357 | 7.8051 |

2. **Covariate**: Age

   **Omics data**: metabolome

Table B.2: Average RMSEs on testing data for predicting Age based on metabolome data and metabolome embeddings from MB-SupCon-cont using three weighting methods (T2D study)

| | **Original** | **MB-SupCon-cont** | | |
|---|---|---|---|---|
| | | Embedding: Linear | Embedding: Exponential | Embedding: Negative-log |
| **ElasticNet** | 6.3917 | 4.6634 | 6.4838 | 5.5157 |
| **SVR** | 8.8224 | 4.4849 | 5.2813 | 5.2183 |
| **RF** | 5.6916 | 4.0593 | 5.0115 | 5.3216 |
| **XGBoost** | 5.7289 | 4.1297 | 4.9475 | 4.8329 |

3. **Covariate**: BMI

   **Omics data**: microbiome

Table B.3: Average RMSEs on testing data for predicting BMI based on microbiome data and microbiome embeddings from MB-SupCon-cont using three weighting methods (T2D study)

| | **Original** | **MB-SupCon-cont** | | |
|---|---|---|---|---|
| | | Embedding: Linear | Embedding: Exponential | Embedding: Negative-log |
| **ElasticNet** | 3.8306 | 3.1463 | 3.3078 | 3.2675 |
| **SVR** | 3.5244 | 3.0928 | 3.1288 | 3.0442 |
| **RF** | 2.9652 | 3.1830 | 3.5324 | 3.4105 |
| **XGBoost** | 3.1530 | 3.1976 | 3.5134 | 3.3364 |

4. **Covariate**: BMI

   **Omics data**: metabolome

Table B.4: Average RMSEs on testing data for predicting BMI based on metabolome data and metabolome embeddings from MB-SupCon-cont using three weighting methods (T2D study)

| | **Original** | **MB-SupCon-cont** | | |
|---|---|---|---|---|
| | | Embedding: Linear | Embedding: Exponential | Embedding: Negative-log |
| **ElasticNet** | 2.8708 | 1.9137 | 2.3156 | 2.0933 |
| **SVR** | 2.8792 | 1.4999 | 1.5844 | 1.6053 |
| **RF** | 2.1848 | 1.4826 | 1.7599 | 1.6190 |
| **XGBoost** | 2.1154 | 1.4799 | 1.8046 | 1.6063 |

5. **Covariate**: SSPG

   **Omics data**: microbiome

Table B.5: Average RMSEs on testing data for predicting SSPG based on microbiome data and microbiome embeddings from MB-SupCon-cont using three weighting methods (T2D study)

| | **Original** | **MB-SupCon-cont** | | |
|---|---|---|---|---|
| | | Embedding: Linear | Embedding: Exponential | Embedding: Negative-log |
| **ElasticNet** | 48.1111 | 44.7639 | 50.9281 | 46.2541 |
| **SVR** | 59.0257 | 44.1470 | 52.6266 | 44.2847 |
| **RF** | 41.9548 | 45.0804 | 54.2055 | 45.4351 |
| **XGBoost** | 44.1595 | 45.5829 | 56.7775 | 46.2306 |

6. **Covariate**: SSPG

**Omics data**: metabolome

Table B.6: Average RMSEs on testing data for predicting SSPG based on metabolome data and metabolome embeddings from MB-SupCon-cont using three weighting methods (T2D study)

| | **Original** | **MB-SupCon-cont** | | |
|---|---|---|---|---|
| | | Embedding: Linear | Embedding: Exponential | Embedding: Negative-log |
| **ElasticNet** | 41.7313 | 22.7196 | 44.2468 | 26.6745 |
| **SVR** | 59.3345 | 24.1464 | 46.7863 | 25.0252 |
| **RF** | 34.4950 | 18.4474 | 46.9629 | 23.4287 |
| **XGBoost** | 33.5242 | 19.1365 | 51.9646 | 23.9354 |

### B.2. Prediction results for HFD study

In HFD study, the mean RMSEs for predictions based on original omics data and on MB-SupCon-cont embeddings are listed here. The prediction results are summarized for covariates Age, BMI and SSPG; omics data of microbiome and metabolome; and predictors ElasticNet, SVR, RF and XGBoost. The dimension of embedding is set to be 10.

1. **Covariate**: Age

   **Omics data**: microbiome

Table B.7: Average RMSEs on testing data for predicting Age based on microbiome data and microbiome embeddings from MB-SupCon-cont using three weighting methods (HFD study)

| | **Original** | **MB-SupCon-cont** | | |
|---|---|---|---|---|
| | | Embedding: Linear | Embedding: Exponential | Embedding: Negative-log |
| **ElasticNet** | 8.6952 | 8.2329 | 9.7006 | 8.4528 |
| **SVR** | 12.5909 | 7.5460 | 10.1288 | 8.2508 |
| **RF** | 8.3439 | 7.8090 | 11.3919 | 8.3139 |
| **XGBoost** | 8.4767 | 7.9057 | 12.3937 | 8.3653 |

2. **Covariate**: Age

   **Omics data**: metabolome

Table B.8: Average RMSEs on testing data for predicting Age based on metabolome data and metabolome embeddings from MB-SupCon-cont using three weighting methods (HFD study)

| | **Original** | **MB-SupCon-cont** | | |
|---|---|---|---|---|
| | | Embedding: Linear | Embedding: Exponential | Embedding: Negative-log |
| **ElasticNet** | 8.9193 | 7.7806 | 9.6889 | 7.7958 |
| **SVR** | 12.8361 | 7.1796 | 10.1834 | 7.7605 |
| **RF** | 9.1652 | 7.1592 | 11.8144 | 7.7460 |
| **XGBoost** | 9.3198 | 7.1434 | 12.7121 | 7.9773 |

3. **Covariate**: Weight

   **Omics data**: microbiome

Table B.9: Average RMSEs on testing data for predicting Weight based on microbiome data and microbiome embeddings from MB-SupCon-cont using three weighting methods (HFD study)

| | **Original** | **MB-SupCon-cont** | | |
|---|---|---|---|---|
| | | Embedding: Linear | Embedding: Exponential | Embedding: Negative-log |
| **ElasticNet** | 8.8105 | 8.3062 | 8.8779 | 8.5224 |
| **SVR** | 10.5297 | 7.9419 | 8.6733 | 8.2047 |
| **RF** | 7.5095 | 8.1008 | 9.0413 | 8.5323 |
| **XGBoost** | 7.8293 | 8.3839 | 9.8254 | 8.6016 |

4. **Covariate**: Weight

   **Omics data**: metabolome

Table B.10: Average RMSEs on testing data for predicting Weight based on metabolome data and metabolome embeddings from MB-SupCon-cont using three weighting methods (HFD study)

| | **Original** | **MB-SupCon-cont** | | |
|---|---|---|---|---|
| | | Embedding: Linear | Embedding: Exponential | Embedding: Negative-log |
| **ElasticNet** | 9.3766 | 7.1195 | 7.8240 | 7.1902 |
| **SVR** | 9.7355 | 6.4042 | 7.6244 | 6.8350 |
| **RF** | 6.9497 | 6.4430 | 7.9505 | 6.8123 |
| **XGBoost** | 7.2810 | 6.4653 | 8.5862 | 6.9363 |
